# An alternatively spliced TREM2 isoform lacking the ligand binding domain is expressed in human brain

**DOI:** 10.1101/2021.11.23.469712

**Authors:** Benjamin C. Shaw, Henry C. Snider, Andrew K. Turner, Diana J. Zajac, James F. Simpson, Steven Estus

**Author notes:** These authors contributed equally to this work (Benjamin C. Shaw, Henry C. Snider). **Correspondence to:** Steve Estus, PhD, Sanders-Brown Center on Aging, Room 537, 789 S. Limestone St, Lexington, KY 40536.

## Abstract

**Background:** Genetic variants in *TREM2* are strongly associated with Alzheimer’s Disease (AD) risk but alternative splicing in *TREM2* transcripts has not been comprehensively described.

**Objective:** Recognizing that alternative splice variants can result in reduced gene expression and/or altered function, we sought to fully characterize splice variation in *TREM2*.

**Methods:** Human blood and anterior cingulate autopsy tissue from 61 donors were used for end-point and quantitative PCR and Western blotting to identify and quantify novel *TREM2* isoforms.

**Results:** In addition to previously described transcripts lacking exon 3 or exon 4, or retaining part of intron 3, we identified novel isoforms lacking exon 2, along with isoforms lacking multiple exons. Isoforms lacking exon 2 were predominant at approximately 10% of TREM2 mRNA in the brain. Expression of *TREM2* and frequency of exon 2 skipping did not differ between AD samples and non-AD controls (p = 0.1268 and p = 0.4909, respectively). Further, these novel splice isoforms were also observed across multiple tissues with similar frequency (range 5.3 – 13.0%). We found that the exon 2 skipped isoform *D2-TREM2* is translated to protein and localizes similarly to full-length TREM2 protein, that both proteins are primarily retained in the Golgi complex, and that D2-TREM2 is expressed in AD and non-AD brain.

**Conclusion:** Since the TREM2 ligand binding domain is encoded by exon 2, and skipping this exon retains reading frame while conserving localization, we hypothesize that *D2-TREM2* acts as an inhibitor of *TREM2* and targeting *TREM2* splicing may be a novel therapeutic pathway for AD.

## Introduction

TREM2 is an activating receptor expressed on innate immune cells, and genetic variants in the *TREM2* gene are associated with both Nasu-Hakola disease and Alzheimer’s Disease (AD) [1–3]. Genome-wide association studies (GWAS) have confirmed that variants rs75932628 and rs143332484, encoding the p.R47H and p.R62H variants, respectively, in *TREM2* are strong risk factors for developing late-onset AD (LOAD) [4–7]. These variants reduce TREM2 function [8–11]. TREM2 is a receptor for both Apolipoprotein E (ApoE) [12] and amyloid beta (Aβ) [13], and regulates Aβ phagocytosis [10, 13, 14], transcriptional changes [15], and microglial transition to a full disease-associated phenotype [16]. Murine models of AD suggest TREM2 may be beneficial early in the disease but detrimental later; *Trem2^KO^* or *Trem2^R47H^* mice crossed with amyloid-based AD models (APP/PS1 or 5xFAD mice) develop greater Aβ pathology [17, 18], but when crossed with the PS19 human tau model, reduced tau pathology [9, 19].

*TREM2* is a five-exon gene that has been reported to undergo alternative splicing [20–22], wherein exon 3 (*D3-TREM2*) or exon 4 (*D4-TREM2*) are skipped, or exon 4 is extended to include a 3’ portion of intron 3. Each of these three alternative splicing isoforms results in a frameshift mutation; exon 3 skipped has been associated with Nasu-Hakola disease as well [21, 22]. Interestingly, though both isoforms encode proteins which lack a transmembrane domain and are expected to be secreted, the exon 4 variants have not been reported as associated with Nasu-Hakola disease possibly due to the lack of an identified causal genetic variant for this isoform. Modulation of *TREM2* splicing has been proposed as a potential therapeutic for Nasu-Hakola disease previously [21], similar to the recent successes with spinal muscular atrophy [23]. Further TREM2 splice variants had not been reported until very recently [24], when an isoform lacking exon 2 (*D2-TREM2),* which encodes the ligand binding domain, was identified. This *D2-TREM2* isoform maintains the reading frame and transmembrane domain but lacks the ligand binding domain.

In this study, we sought to fully characterize *TREM2* alternative splicing in brain. We identify many more alternative splice isoforms than previously reported, and report that this alternative splicing is not brain-specific as it is conserved across multiple tissues. Further, we show that the *D2-TREM2* splice isoform is translated into protein using overexpression paradigms, and that this D2-TREM2 protein has similar compartment localization as the fulllength (FL-TREM2) protein. We propose that modulating *D2-TREM2* could be exploited to enhance TREM2 function—by decreasing *D2-TREM2* early, one could increase the then-beneficial functional TREM2; by increasing *D2-TREM2* late in disease, one could inhibit the then-detrimental functional TREM2.

## Materials and Methods

### Preparation of DNA, RNA, cDNA, and protein from human samples

Human blood and anterior cingulate autopsy tissue from 61 donors were generously provided by the Sanders-Brown Alzheimer’s Disease Center neuropathology core and their characteristics and cDNA synthesis have been described elsewhere [25]. All human subjects research was carried out in accordance with the University of Kentucky Institutional Review Board under protocol number 48095. The matched brain and blood samples were from deceased individuals with an average age at death of 82.4 ± 8.7 (mean ± SD) years for non-AD and 81.7 ± 6.2 years for AD subjects. The average postmortem interval (PMI) for non-AD and AD subjects was 2.8 ± 0.8 and 3.4 ± 0.6 hrs, respectively. Non-AD and AD samples were comprised of 48% and 55% female subjects, respectively. MMSE scores were, on average, 28.4 ± 1.6 for non-AD subjects and 11.9 ± 8.0 for AD subjects. Total RNA was prepared using a Qiagen RNeasy Lipid Tissue Mini kit (Qiagen #74804) according to manufacturer’s instructions. Reverse transcription was carried out using SuperScript IV (Invitrogen #18091050) according to manufacturer’s instructions. For cross-tissue splicing comparison, fetal RNA libraries from various tissues were obtained from a commercial vendor (Stratagene) and their cDNA preparation has been described elsewhere [26]. Protein homogenates were obtained from hippocampal samples of three AD and three non-AD individuals. Approximately 100 mg of tissue was added to two volumes of sucrose buffer (0.25 M sucrose, 20 mM EDTA, 20 mM EGTA, 100 mM Tris, pH 7.4) and homogenized with a pestle in a microfuge tube for 25 strokes. After homogenization, one volume of a 2X solution of RIPA buffer (300 mM NaCl, 2% NP-40, 2% w/v deoxycholate, 0.2% sodium dodecyl sulfate, 50 mM Tris-HCl, pH 7.4) was added for a final 1X RIPA solution. Both the sucrose and 2X RIPA buffers contained a 1X final concentration of protease and phosphatase cocktail (Invitrogen #78442).

### Cell Culture

The HMC3 human microglial cell line was obtained from American Type Culture Collection (ATCC CRL-3304). HEK293 and HMC3 cells were cultured in Eagle’s Modified Minimum Essential Medium (EMEM), ATCC modification (ATCC 30-2003) supplemented with 10% fetal bovine serum, defined (HyClone, GE Healthcare SH30070.03); 50 U/mL penicillin, 50 μg/mL streptomycin (Gibco 22400-089). Cells were maintained at 37°C in a 5% CO2 in air atmosphere.

### TREM2 splice isoform identification by PCR and sequencing

The cDNA samples from the anterior cingulate samples were amplified using primers corresponding to *TREM2* exon 1 (5’-CCTGACATGCCTGATCCTCT-3’) and exon 5 (5’-GTGTTCTTACCACCTCCCC-3’) with Q5 high-fidelity hot-start polymerase (NEB # M0493L). Thermocylcing parameters were as follows: 98°C 30 s; 98°C 5 s, 67°C 5 s, 72°C 45 s, 30 cycles; 72°C 2 min, 25°C hold. PCR products were separated on a 8% acrylamide gel and imaged using a BioRad ChemiDoc XR. Bands were extracted for subsequent amplification as above and purification using a Monarch PCR Cleanup Kit (NEB T1030L). Purified products were sequenced commercially (ACGT; Wheeling, IL) and compared to the reference transcript NM_018965.4 to determine splicing.

### Quantifcation of TREM2 transcripts

Quantitative PCR (qPCR) was used to quantify expression of *TREM2* transcripts. Primers corresponding to sequences within exons 1 and 2 were used to quantify *TREM2* exon 2 expression (forward, 5’-CCTTGGCTGGGGAAGGG-3’; reverse, 5’-TCATAGGGGCAAGACACCTG-3’), as well as primers corresponding to sequences at the exon 1–3 junction and within exon 3 to quantify the *D2-TREM2* isoform (forward, 5’-TTACTCTTTGTCACAGACCCC-3’; reverse, 5’-GGGCATCCTCGAAGCTCT-3’). PCR was conducted using an initial 2 min incubation at 95°, followed by 40 cycles of 10 s at 95°C, 20 s at 60°C, and 20 s at 72°C. The 20 μl reactions contained 1 μM each primer, 1X PerfeCTa SYBR Green Super Mix (Quanta Biosciences), and 20 ng of cDNA. Experimental samples were amplified in parallel with serially diluted standards that were generated by PCR of cDNA using the indicated primers, followed by purification and quantitation by UV absorbance. Results from samples were compared relative to the standard curve to calculate copy number in each sample. Total TREM2 expression was the sum of the copy numbers for TREM2 exon 2 present and exon 2 skipped. Assays were performed in duplicate and normalized to expression of Iba1 (*AIF1*) as the housekeeping gene, as *TREM2* in the CNS is exclusively expressed in microglia. For cross-tissue comparison, the percent exon 2 skipping was calculated by dividing the exon 2 skipped copies by the sum of the exon 2 skipped and mean of exon 2 present copies without normalization. Data in the cross-tissue comparison reflect six technical replicates.

### TREM2 transcript cloning

For immunofluorescence experiments, cDNA corresponding to the full-length *TREM2* and *D2-TREM2* transcripts was PCR-amplified using the primers corresponding to sequence within exons 1 and 5, described above. For a D2-TREM2 size standard for Western blotting, the *D2-TREM2* transcript was cloned with the same exon 1 primer and with the reverse primer corresponding to sequence in the 3’UTR (5’–CCAGCTAAATATGACAGTCTTGGA – 3’) to preclude an epitope tag. Amplification was performed with Platinum Taq (Invitrogen 10966034) with the following cycling parameters: 2 min at 94°C; 30 s at 94°C, 30 s at 60°C, 2 min at 72°C, 30 cycles; 7 min at 72°C, 25°C hold. All cloning was performed using a pcDNA 3.1-V5/His TOPO-TA cloning kit (Invitrogen K480001) per manufacturer’s instructions. Clones were verified by sequencing (ACGT; Wheeling, IL) and grown for midi-scale production and purification using a Qiagen Plasmid Plus Midiprep kit (Qiagen 12943).

### HMC3 and HEK293 Transfection

HMC3 human microglia or HEK293 cells were transfected with Lipofectamine 3000 with Plus reagent (Invitrogen L3000001) per manufacturer instructions with 0.8 μL of Lipofectamine 3000, 1 μL Plus reagent, and 250 ng plasmid per well in 8 well glass chamber slides (MatTek CCS-8). Cells were incubated for 24 hours prior to processing for microscopy (HMC3) or Western blotting (HEK293).

### Confocal Immunofluorescence Microscopy

Transfected HMC3 cells were fixed with 10% neutral buffered formalin (Fisher Scientific SF100-4) for 30 minutes then blocked and permeabilized for 30 minutes with 10% goat serum (Sigma S26-LITER), 0.1% Triton X-100 (Fisher Scientific BP151-500) in PBS (Fisher BioReagents BP665-1). Primary and secondary antibodies were diluted in the same blocking and permeabilization buffer and incubated at room temperature for 90 minutes. Cells were washed three times in blocking and permeabilization buffer between primary and secondary antibodies, and three times in PBS prior to coverslip mounting with Prolong Glass with NucBlue mounting media (Invitrogen P36981) and high-tolerance No. 1.5 coverglass (ThorLabs CG15KH1). Images were acquired using a Nikon A1R HD inverted confocal microscope with a 60X oil objective and NIS Elements AR software.

### Western Blotting

Protein was quantified using a Pierce 660 assay (Invitrogen #1861426) and 40 μg loaded per well on a 10-20% Tricine gel (Invitrogen #EC6625BOX) followed by transfer to a 0.22 μm PVDF membrane. The membrane was blotted overnight with an antibody against sequence within the TREM2 cytosolic domain (Cell Signaling #91068) and probed with a goat anti-rabbit secondary AlexaFluor 800 secondary antibody (Invitrogen #A32735).

### Statistical Analyses

Analyses were performed using GraphPad Prism 8.4.2. Quantitative data were first checked for normality by the D’Agostino & Pearson test. Normally distributed data were analyzed using a two-tailed t-test, while data not normally distributed were analyzed with a twotailed Mann Whitney test. The type of test is noted along with p values.

## Results

### TREM2 undergoes extensive alternative splicing in human adult brain

To fully characterize *TREM2* alternative splicing, we PCR-amplified *TREM2* cDNA from anterior cingulate cortex by using primers corresponding to sequences within exons 1 and 5. A gene map, including introns and encoded protein domains, is shown with forward and reverse primers (Figure 1, top). We observed substantial alternative splicing in both AD and non-AD individuals (Figure 1, left). The identity of each isoform was confirmed by direct sequencing (Figure 1, right). This effort identified multiple novel TREM2 isoforms that have not been reported previously, including isoforms with multiple exons skipped.

**Figure 1:**
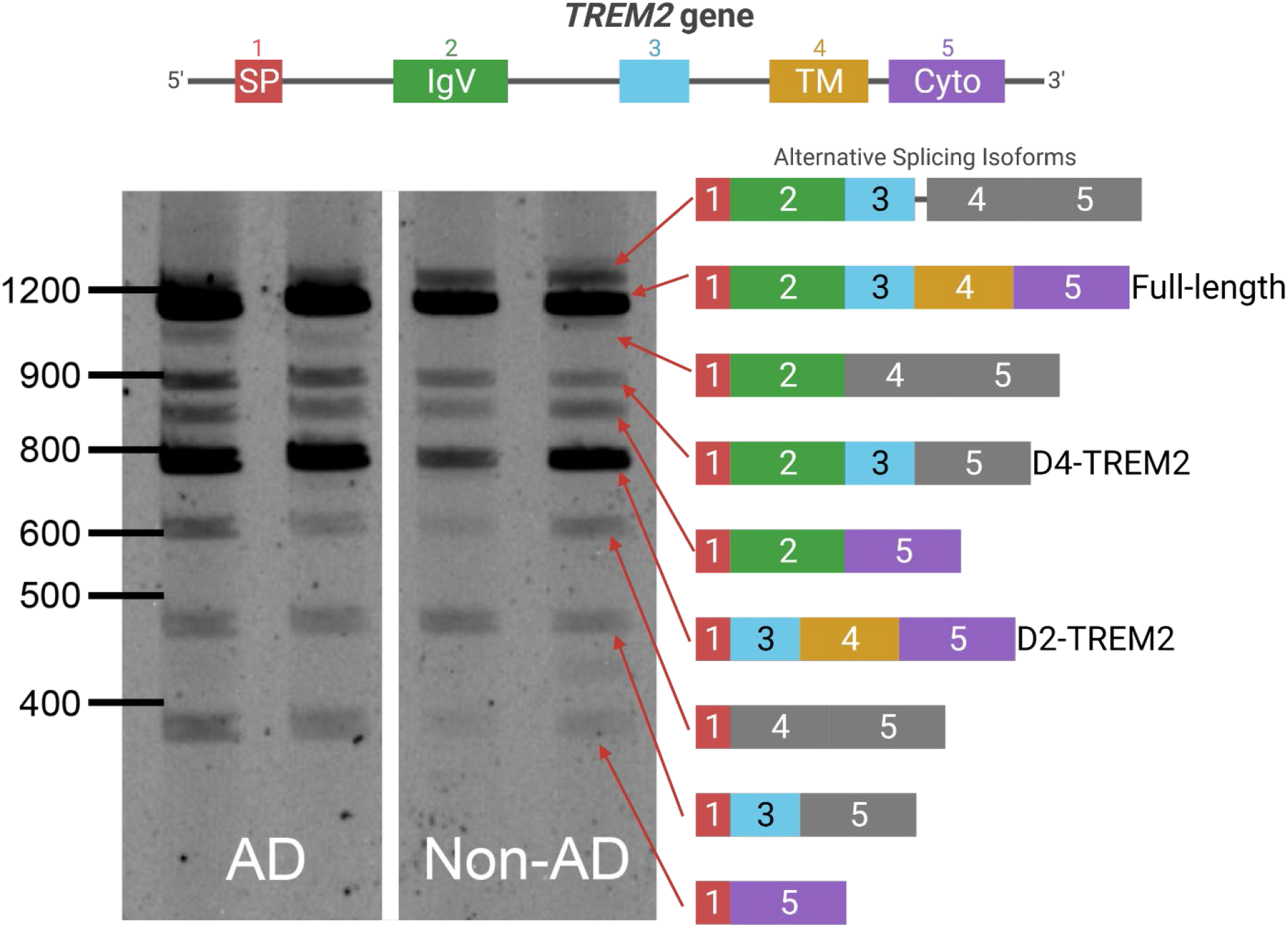
*TREM2* undergoes extensive alternative splicing. Top: a schematic of the *TREM2* gene is shown with introns and exons. Left: a representative image of the PCR amplification and gel electrophoresis of *TREM2* cDNA from AD and non-AD brains. No differences in splicing patterns were noticed between AD and non-AD. Right: schematic of the splice isoforms identified after sequencing. Colors correspond to exons in the gene model, while frameshifts are shown in grey. A doublet appears in the lower transcripts on the gel in which the lower band corresponds to *CES3* in addition to the identified *TREM2* isoforms. All bands identified in Figure 1 have been confirmed by Sanger sequencing.

Since this assay suggested that the isoform lacking exon 2 (*D2-TREM2*) is the most abundance variant, we next quantified *TREM2* and this alternate *D2-TREM2* isoform in a series of brain samples. Since *TREM2* is almost exclusively expressed by microglia in the brain, we normalized the *TREM2* copy number to that of *AIF1* (Figure 2A). While total *TREM2* expression is not significantly different with respect to AD (p = 0.1268, two-tailed t-test), there is a significant increase as a function of high vs. low pathology (NIARI = 3 vs. NIARI < 3; Figure 2B; p = 0.0014, Welch’s t-test). We next investigated whether *D2-TREM2* occurred more frequently in AD vs. non-AD individuals. We found that exon 2 skipping correlates well with total TREM2 expression (Figure 2C), but observed no difference in exon 2 skipping frequency between the two groups (Figure 2D; p = 0.4909, Mann Whitney test).

**Figure 2:**
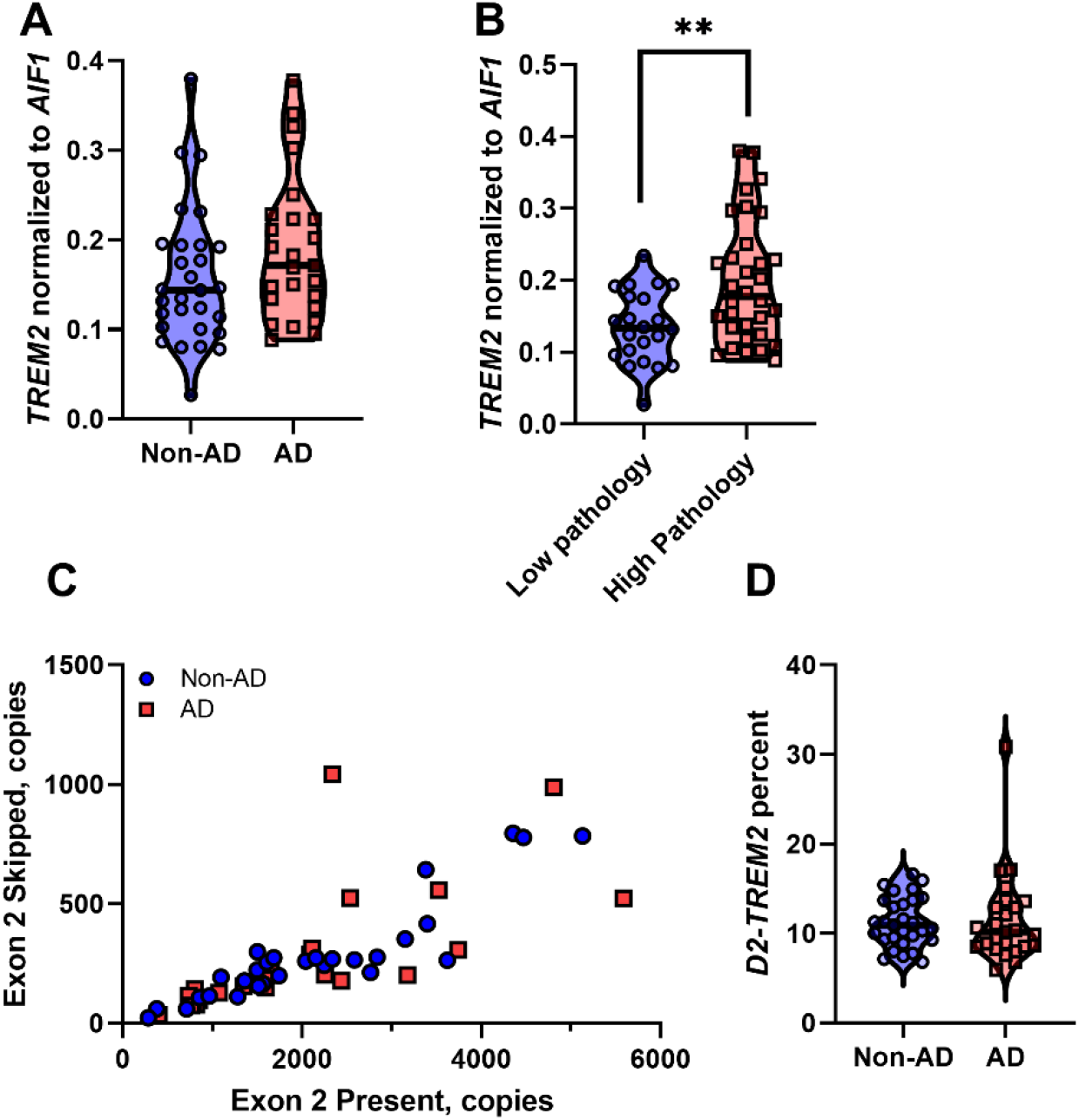
Quantification of *TREM2* and exon 2 skipping in human brain tissue. (A): Total *TREM2* expression normalized to the microglial marker *AIF1* expression. *TREM2* expression is not significantly different between AD and non-AD samples (p = 0.1268, t-test). (B) Total *TREM2* expression normalized to *AIF1* is increased when parsed by high pathology (NIARI = 3) vs. low pathology (NIARI > 3), (p = 0.0014, Welch’s t-test). (C) Expression of the isoform lacking exon 2 correlates well with expression of the isoform containing exon 2. (D) Exon 2 is skipped at an approximate frequency of 11%, with no significant difference between AD vs. non-AD (p = 0.4909, Mann-Whitney test).

### TREM2 alternative splicing is conserved across tissues

To test whether the observed alternative splicing is specific to brain, we subjected cDNA libraries from aorta, lung, kidney, heart, skeletal muscle, brain, and liver to PCR amplification with primers in TREM2 exons 1 and 5, as described above (Figure 1). We observed each of the previous splice isoforms across multiple tissues, though each isoform was not present in all tissues (Figure 3). We also observed a high relative abundance of the isoform lacking both exon 2 and 3 along with the isoform lacking exons 2 and 4. This may reflect PCR bias, where shorter fragments are amplified more efficiently than longer fragments and are overrepresented in relative abundance. Nonetheless, the *D2-TREM2* isoform was an abundant alternative isoform across all tissues surveyed. We then quantified *TREM2* exon 2 skipping frequency in these samples and found similar frequency of exon 2 skipping across tissues (Figure 4).

**Figure 3:**
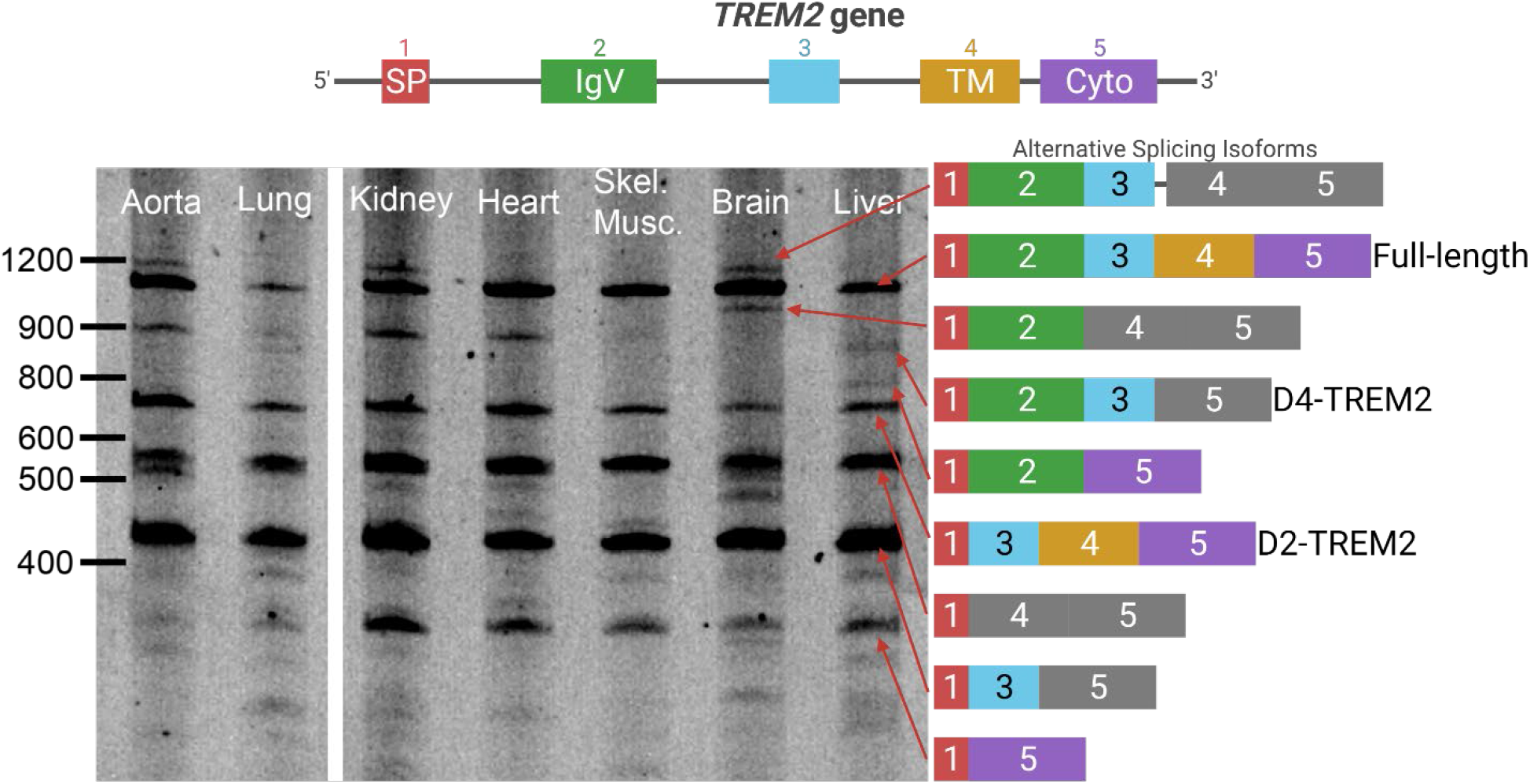
Complex patterns of *TREM2* alternative splicing are present in many tissues. Human fetal cDNA libraries from multiple tissues were amplified using the same primers from Figure 1. The splice variants from Figure 1 are replicated across these six additional tissues. Differences in the relative abundance of each splice variant may be due to differences in developmental stage and/or splicing factor differences between tissues.

**Figure 4:**
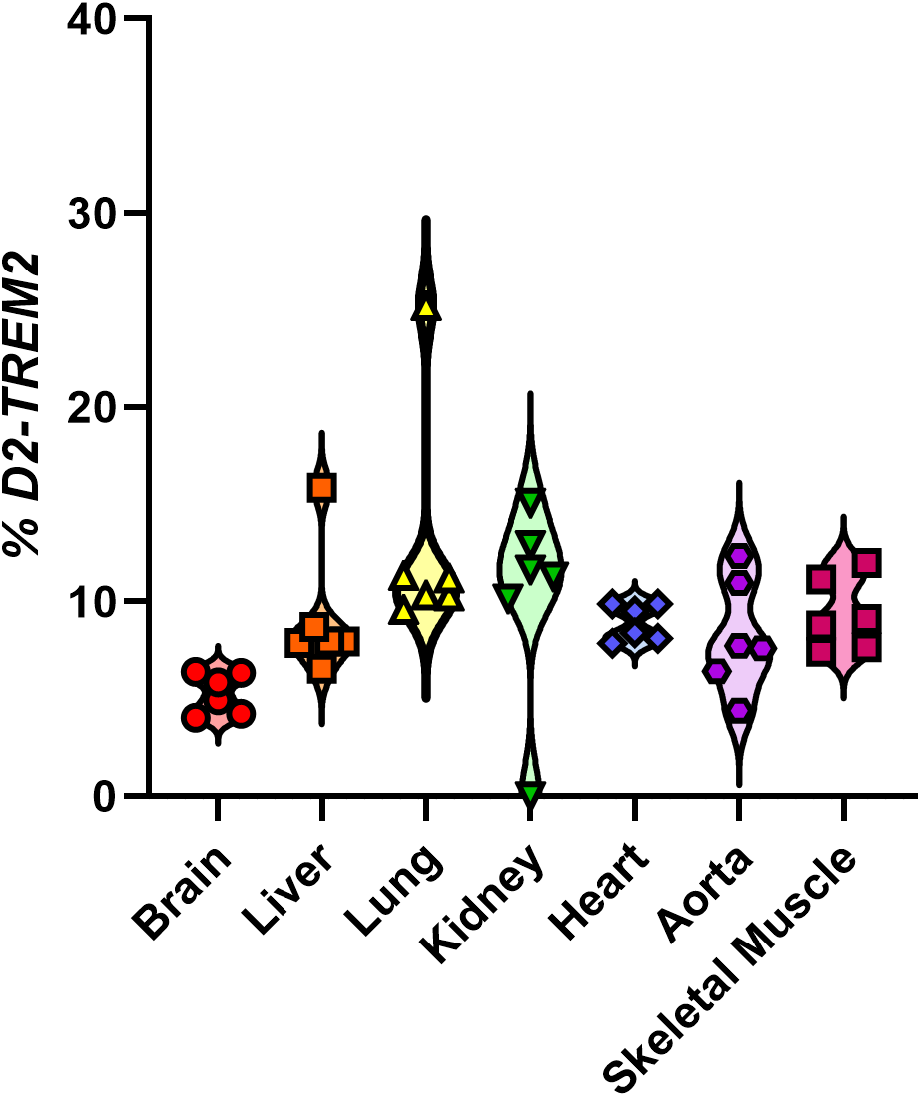
Quantification of *TREM2* and exon 2 skipping across tissues. Exon 2 is skipped at a frequency between 5.30 – 13.0%. Data points reflect technical, not biological, replicates from pooled cDNA libraries.

### D2-TREM2 protein localizes similar to full-length TREM2

To test whether the D2-TREM2 protein isoform is trafficked similarly to the full-length isoform, we cloned each isoform into expression vectors and transfected HMC3 human microglial cells for confocal microscopy (Figure 5). We observed similar staining patterns from both the *FL-TREM2* (Figure 5A) and *D2-TREM2* (Figure 5B), and this staining pattern is consistent with intracellular retention prevoiusly reported [27, 28]. This implies the D2-TREM2 protein maintains the localization pattern of its parent full-length TREM2. We confirmed the D2-TREM2 isoform is predominantly retained in the Golgi complex (Figure 5C) as has been previously reported for the FL-TREM2 and shown here (Figure 5D).

**Figure 5:**
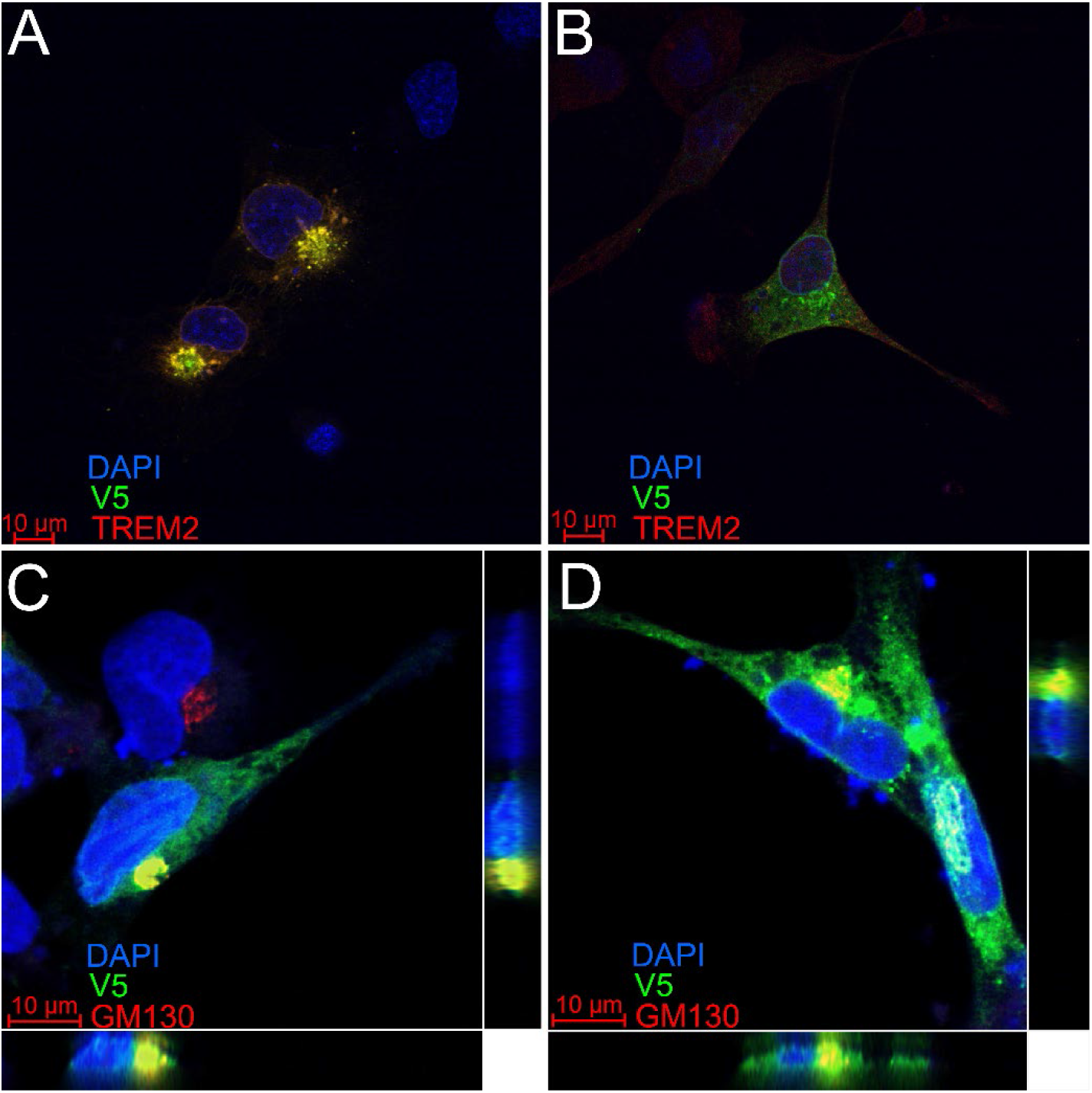
TREM2 and D2-TREM2 have a similar subcellular localization. The HMC3 human microglial cell line was transfected with vectors encoding either *full-length TREM2* (A,C) or *D2-TREM2* (B,D). Both vectors have an in-frame V5 epitope tag. In A and B, cells were subsequently labeled with antibodies against a TREM2 epitope encoded by exon 2 (red) or V5 (green). The limited red fluorescent labeling and absence of yellow overlap in the D2-TREM2 transfected cells (B) is due to low endogenous TREM2 expression. In C and D, transfected cells were labeled with antibodies against the Golgi complex marker GM130 (red) and V5 (green). The intracellular pools of both TREM2 (C) and D2-TREM2 (D) largely colocalize with GM130. Views represent the XY (main), XZ (bottom) and YZ (right) views in each panel.

### D2-TREM2 protein is observed in human hippocampus samples

To test whether the protein encoded by the *D2-TREM2* transcript is present in human brain, hippocampal protein from three AD and three non-AD individuals was subjected to Western blotting with an antibody that recognizes the cytosolic tail of TREM2 (Figure 6). A short exposure (Figure 6A) showed the previously described immature, non-glycosylated and mature, glycosylated full-length TREM2 as well as the carboxyl-terminal TREM2 fragment. A size reference for D2-TREM2 was included on the blot using HEK293 cells transfected with a *D2-TREM2* cDNA vector. A very short exposure shows that two proteins are labeled by the TREM2 antibody in the transfected cells, but not in the non-transfected cells, confirming these bands reflect products of *D2-TREM2*. A longer exposure (Figure 6B) reveals a doublet of bands in human brain at 12 and 13 kDa which matches the pattern observed in the *D2-TREM2* transfected cells. Considering these apparent molecular weights, we note that the carboxyl-terminal TREM2 fragment has a predicted molecular weight of 8 kDa while the predicted molecular weight of D2-TREM2 is 11 kDa. Hence, both the carboxyl-terminal fragment and D2-TREM2 migrate 3 kDa heavier than predicted. In summary, D2-TREM2 appears present in both AD and non-AD human brain.

**Figure 6:**
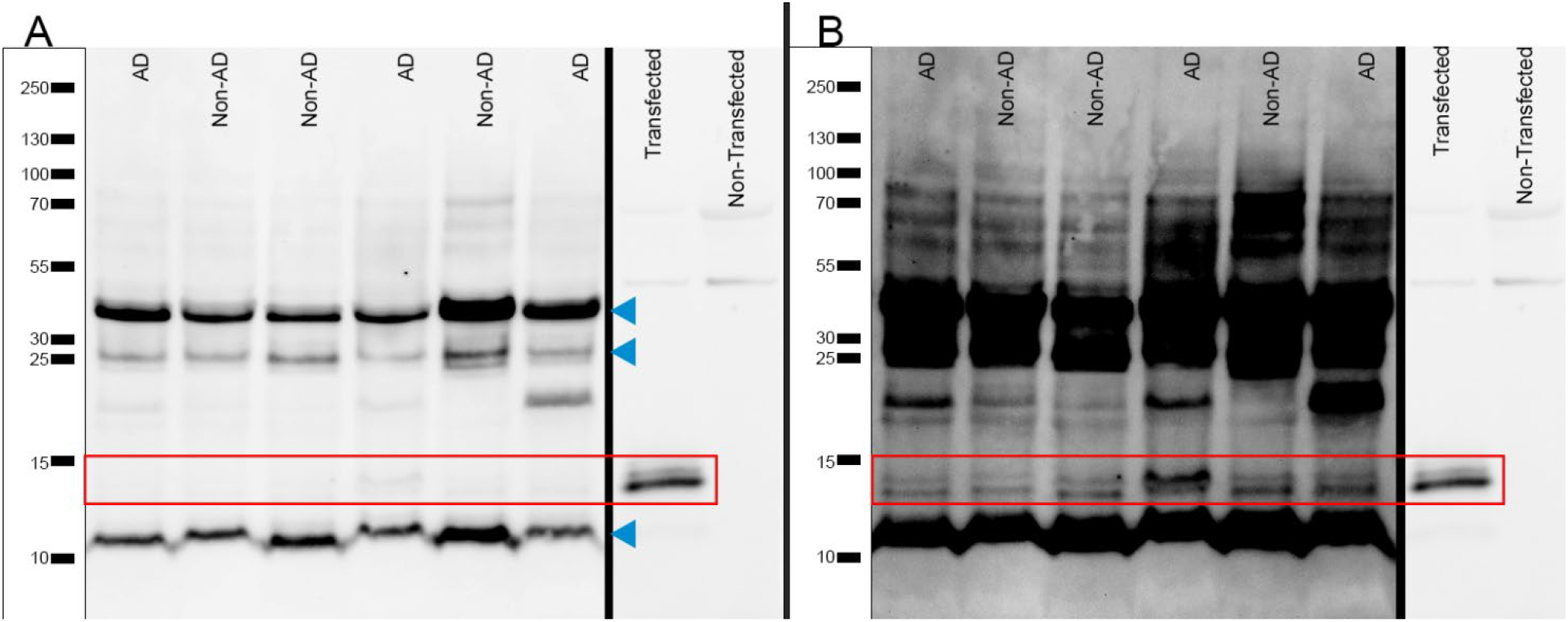
Western blot of human brain detects D2-TREM2 protein. Full-length glycosylated and immature TREM2 protein, and its proteolytic carboxyl-terminal fragment are marked with blue triangles. The *D2-TREM2* protein products migrate at 12 and 13 kDa (red box), slightly higher than the carboxyl-terminal fragment. The protein is faintly visible with a lower exposure (A), but more robust with longer exposure (B). D2-TREM2 transfected HEK293 cells were used to generate a size standard in these experiments and blot images were taken at a reduced exposure that is included alongside both images to demonstrate that the protein doublet corresponding to D2-TREM2 in cell culture matches the doublet observed in brain.

## Discussion

The primary findings of this report are that *TREM2* undergoes far more extensive alternative splicing than has been previously reported [20–22, 24], that D2-TREM2 is a common variant that is not influenced by AD neuropathology, and that D2-TREM2 co-localizes with fulllength TREM2. As such, this comprehensive analysis of *TREM2* alternative splicing extends prior knowledge, noting that Ensembl lists only full-length TREM2 (ENST00000373113.8), exon 4 skipped (*D4-TREM2;* ENST00000338469.3), and intron 3 retained isoforms (ENST00000373122.8) as known transcripts; NCBI only lists the full length (NM_018965.4) and *D4-TREM2* (NM_001271821.2). Exon 3 skipping has been previously described in two small cohorts [3, 22]. The *D2-TREM2* isoform is not annotated in either the NCBI nor Ensembl database. The *D2-TREM2* isoform in brain was recently reported [24, 29], and our brain quantitation data closely replicate their results (Figure 2C). In addition, we demonstrate that this splice isoform is expressed across multiple tissues (Figure 3) with roughly equal relative abundance (Figure 4). We further extend these recent findings by demonstrating that isoforms exist with multiple exons skipped, such that all permutations of exons 2, 3, and 4 can be skipped individually or in combination (Figure 1), and that exon 2 skipping frequency is not different as a function of AD neuropathology.

While TREM2 has well-described cell surface localization including receptor activity [12–14, 17], interactions with DAP12 [30], and proteolytic cleavage by ADAM10 [31, 32], there also exists a considerable intracellular TREM2 pool [27, 28]. Our localization studies support the hypothesis that D2-TREM2 localizes to the same compartment as FL-TREM2. Both appear to be predominantly localized to the Golgi complex (Figure 5C-D), replicating previous work with FL-TREM2 [27, 28]. The recent *D2-TREM2* report [24] provided evidence of membrane-bound D2-TREM2 and FL-TREM2 by Western blots on subcellular fractions. Taken together, this suggests that TREM2—and likely D2-TREM2—resides in a transmembrane Golgi pool. We are also the first to show that the *D2-TREM2* isoform is present as protein in human brain (Figure 6). While the *D2-TREM2* transcript comprises approximately 10% of the total *TREM2* mRNA, the protein appears less abundant than the mRNA would predict. We speculate that this may be due to reduced translation efficiency or protein instability.

Functionally, TREM2 seems to operate under a feed-forward mechanism which D2-TREM2 may act to inhibit. Experiments *in vitro* suggest that the transmembrane Golgi pool rapidly moves to the cell membrane in response to intracellular calcium flux induced by ionomycin [27, 28]. Ligation of TREM2 using monoclonal antibodies also elicits an intracellular calcium spike. Whether D2-TREM2 inhibits this feed-forward mechanism, where TREM2 signaling induces calcium flux and increases cell surface mobilization, is still unknown.

*TREM2* variants were some of the first genetic risk factors outside of *APOE4* identified for LOAD [1, 2], have been replicated in multiple GWAS [4–7], and have the highest odds ratios for LOAD after *APOE4* [33]. Variants encoding the p.R47H (rs75932628) and p.R62H (rs143332484) disrupt ligand binding and are predicted to be partial loss-of-function mutations [8–11]. We hypothesize that the D2-TREM2 protein is a functionally null receptor, as the ligand binding IgV domain is missing (Figure 1), though this has yet to be confirmed. In a similar case, *CD33* has an isoform with relatively high abundance that is also lacking its ligand binding IgV domain (*D2-CD33);* however genetics [34], mouse models [35], and *in vitro* studies with human pluripotent stem cell-derived microglia [36, 37] suggest the D2-CD33 protein may have a gain of function which acts independent of its loss of receptor activity.

*TREM2* remains a promising target in potential AD therapeutics, evidenced by the strong interest in preclinical TREM2 monoclonal antibody treatments including a Phase 2 trial of AL002a (Alector, NCT04592874). Modulation of alternative splice isoforms of *TREM2* may represent a novel therapeutic pathway. The previously reported *D4-TREM2* isoform lacks a transmembrane domain (Figure 1) and is likely secreted as soluble TREM2 (sTREM2) which may be AD-protective [38, 39]. We posit that the *D2-TREM2* isoform acts similarly to the p.R47H and p.R62H variants by reducing TREM2 function, and balanced with *D4-TREM2,* this forms a potentially druggable rheostat. Early in the disease, increased TREM2 function and sTREM2 secretion are apparently beneficial; increasing full-length *TREM2* would promote functional TREM2 signaling and increased *D4-TREM2* would promote increased sTREM2 secretion. Later in disease when inflammation may be detrimental, increasing *D2-TREM2* could provide a switch to decrease TREM2 activity and inflammation (Figure 7).

**Figure 7:**
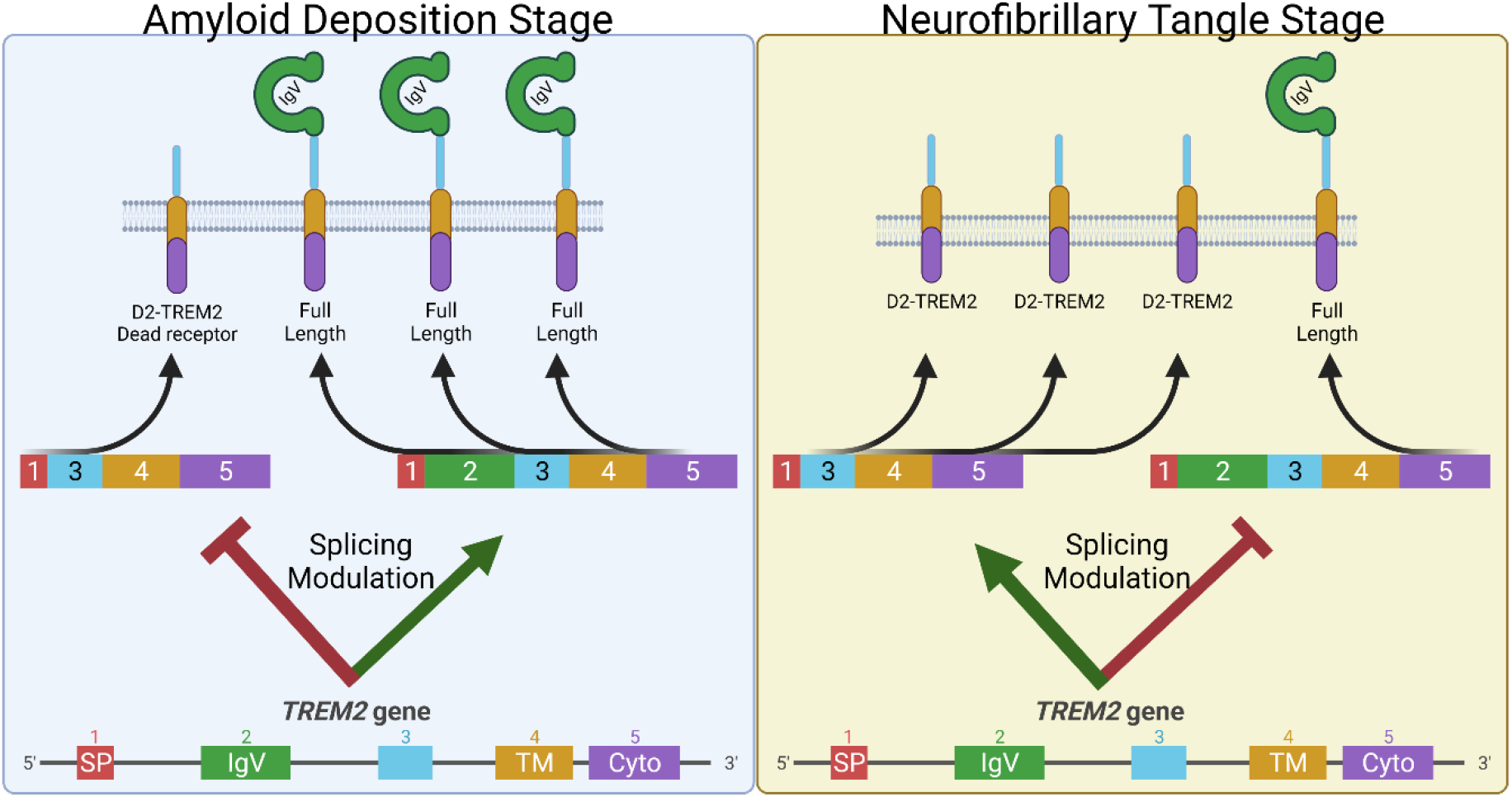
Model to exploit alternative splicing in *TREM2* as a potential AD therapeutic. Early in disease when Aβ pathology is developing, animal models suggest TREM2 signaling is protective. Hence, decreasing *D2-TREM2* and increasing full-length *TREM2* may be helpful in this stage of the disease (Left). Animal models also indicate TREM2 deficiency is protective from tau-related pathology suggesting a detrimental role for TREM2 signaling in this later stage of the disease when hyperphosphorylated tau accumulates. At this point, increased skipping of exon 2 to promote the “dead receptor” D2-TREM2 and thereby inhibit TREM2 signaling may be helpful (Right).

## Acknowledgements

This work was supported by F99NS120365 (BCS), RF1AG059717 (SE), and R21AG068370 (SE). We would like to thank the Sanders-Brown Center on Aging Biorepository for samples, supported by P30AG072946.

## Authorship Contributions

**Benjamin C. Shaw:** Conceptualization, Methodology, Software, Validation, Formal Analysis, Investigation, Data Curation, Writing – Original Draft, Writing – Review & Editing, Visualization, Supervision, Funding Acquisition. **Henry C. Snider:** Conceptualization, Validation, Investigation, Writing – Review & Editing, Visualization. **Andrew K. Turner:** Investigation, Writing – Review & Editing, Visualization. **Diana J. Zajac:** Validation, Investigation, Writing – Review & Editing, Visualization. **James F. Simpson:** Conceptualization, Methodology, Software, Validation, Formal Analysis, Investigation, Data Curation, Writing – Review & Editing, Visualization. **Steven Estus:** Conceptualization, Methodology, Software, Validation, Formal Analysis, Resources, Data Curation, Writing – Review & Editing, Visualization, Supervision, Project Administration, Funding Acquisition.

